# The histone variant H2A.W cooperates with chromatin modifications and linker histone H1 to maintain transcriptional silencing of transposons in *Arabidopsis*

**DOI:** 10.1101/2022.05.31.493688

**Authors:** Pierre Bourguet, Ramesh Yelagandula, Taiko Kim To, Akihisa Osakabe, Archana Alishe, Rita Jui-Hsien Lu, Tetsuji Kakutani, Pao-Yang Chen, Frédéric Berger

**Affiliations:** Gregor Mendel Institute (GMI), Austrian Academy of Sciences, Vienna Biocenter (VBC), Dr. Bohr Gasse 3, 1030 Vienna, Austria; Institute of Molecular Biotechnology of the Austrian Academy of Sciences (IMBA), Vienna Biocenter (VBC), Vienna, Austria; Department of Biological Sciences, The University of Tokyo, Tokyo, Japan; Institute of Plant and Microbial, Academia Sinica, 11529, Taipei, Taiwan

**Keywords:** heterochromatin, histone, histone H2A variants, epigenetics, transcriptional silencing

## Abstract

Transposable elements (TEs) are marked by a complex array of chromatin modifications, but a central unifying mechanism for how they are silenced remains elusive. Histone H3 Lysine 9 methylation (H3K9me) is an important component of heterochromatin in most eukaryotes, including plants. In flowering plants, the specialized histone variant H2A.W occupies nucleosomes found at TE sequences. This variant is deposited by the chromatin remodeler DDM1 and confers specific biophysical properties to the nucleosomes.

Here we use genetic and genomic strategies to evaluate the role of H2A.W in transposon silencing in Arabidopsis. Compared with mutants lacking either H2A.W or H3K9me, the combined loss of both H2A.W and H3K9me causes a dramatic increase in both the number of expressed TEs and their expression levels. Synergistic effects are also observed when H2A.W is lost in combination with histone H1 or CH methylation. Collectively, these TEs are also upregulated in mutants lacking DDM1, which are impaired in H2A.W deposition and lose heterochromatic marks.

We conclude that H2A.W acts in combination with different elements of heterochromatin to maintain silencing across a large spectrum of TEs present primarily in pericentric heterochromatin in Arabidopsis. In mammals, the DDM1 ortholog LSH deposits macroH2A to heterochromatin and silences TEs. We thus propose that specialized H2A variants localized to heterochromatin interact with a complex array of histone modifications to silence TEs in eukaryotes.

## Introduction

Transposable elements (TEs) are DNA sequences able to duplicate and change their position in the genome via a multi-step mechanism called transposition. As transposition requires the expression of TE sequences, host genomes have evolved diverse strategies to prevent transposition using transcriptional and post-transcriptional silencing. Like all components of the eukaryotic genome, TEs are packaged in nucleosomes formed by the assembly of a tetramer of the core histones H3 and H4 with two heterodimers of the core histones H2A and H2B. TEs tend to be segregated within specific chromatin features collectively called heterochromatin (Allshire & Madhani, 2017). In plants and mammals, heterochromatic nucleosomes are characterized by both specific histone modifications and specific histone variants (Martire & Banaszynski, 2020). In mammals, the specialized variant macroH2A tends to be enriched at TEs, although it is also found in other contexts (Sun & Bernstein, 2019). Flowering plants evolved the heterochromatin-specific variant H2A.W, which is exclusively present in nucleosomes at TEs (Kawashima et al., 2015; Talbert & Henikoff, 2021; Yelagandula et al., 2014). While H2A.W confers specific properties to nucleosomes that lead to heterochromatin compaction (Bourguet et al., 2021; Osakabe et al., 2018; Yelagandula et al., 2014), its role in TE silencing remains to be studied in further details.

This is in contrast with histone modifications and DNA methylation in symmetrical (CG, CHG) and non-symmetrical (CHH) contexts, which are well known factors of TE silencing (Du et al., 2015; Rigal & Mathieu, 2011). CHH methylation is partially maintained by the RNA dependent DNA methylation (RdDM) pathway, which relies on the complementarity of non-coding RNAs to TE sequences (Wendte & Pikaard, 2017). The RdDM pathway primarily methylates short TEs and the edges of long TEs, while CHH methylation within TE bodies is maintained by the flowering plant-specific CHROMOMETHYLASE 2 (CMT2) (Stroud et al., 2014; Zemach et al., 2013). CHG methylation is maintained by the land plant-specific CHROMOMETHYLASE 3 which, like CMT2, possesses a chromo domain that binds histone H3 lysine 9 mono or di-methylation (hereafter referred to together as H3K9me) (Du et al., 2012; Stroud et al., 2014). H3K9me also recruits the RdDM pathway (Law et al., 2013; H. Zhang et al., 2013), making it a central hub for DNA methylation. Moreover, H3K9me is bound by Agenet domain (AGD)-containing p1 (AGDP1) to promote chromatin condensation and TE silencing, possibly through DNA methylation (C. Zhang et al., 2018; Zhao et al., 2018). In land plants, the bulk of DNA methylation depends on DNA METHYLTRANSFERASE 1 (MET1), the ortholog of the mammalian *Dnmt1*, which deposits CG methylation and maintains TE silencing in a H3K9me-independent fashion (Du et al., 2015).

Compared to those TEs silenced by H3K9me or CG methylation, an even larger number of TEs are silenced by the chromatin remodeler DECREASED DNA METHYLATION 1 (DDM1) (Oberlin et al., 2017; Osakabe et al., 2021; Stroud et al., 2012). Absence of DDM1 results in loss of DNA methylation in all contexts as well as H3K9me and H3K27me1 (Gendrel et al., 2002; Ikeda et al., 2017; Vongs et al., 1993; Zemach et al., 2013). The loss of DNA methylation in *ddm1* mutants depends on linker histone H1 (Lyons & Zilberman, 2017; Zemach et al., 2013). However, there is no evidence for DDM1 directly affecting H1 dynamics. Instead, it binds to H2A.W through two conserved binding sites and deposits H2A.W at TEs that are expressed in *ddm1* knock-out mutants (Osakabe et al., 2021). The interaction between DDM1 and H2A.W is required for DDM1-mediated TE silencing, leading to the hypothesis that H2A.W plays a direct role in transcriptional TE silencing. However, loss of H2A.W does not result in increased transcription of TEs but instead, when combined with the loss of H1, increases chromatin accessibility to TEs (Bourguet et al., 2021). Therefore, we speculated that H2A.W cooperates with heterochromatin components to silence TEs targeted by DDM1.

Here, we report that H2A.W silences distinct sets of TEs synergistically with H1 or H3K9me. Genetic interactions place H2A.W downstream of DDM1 and analyses of transcriptomes in different genetic backgrounds show that the synergies between H2A.W and either H3K9me or H1 account for half of TEs silenced by DDM1, which represents a quarter (26%) of all TEs in the Arabidopsis genome.

## Results

### H2A.W contributes to TE silencing in conjunction with other heterochromatic marks

While H2A.W is specifically incorporated at transcriptionally silent TEs (Yelagandula et al., 2014), *h2a.w-2* mutations have little effect on TE transcription and do not affect H3K9me (Bourguet et al., 2021). What is observed in these mutants is an increased enrichment of linker histone H1 over pericentromeres (Bourguet et al., 2021). Hence, we reasoned that H1 and H3K9me could act redundantly with H2A.W to silence TEs and could be sufficient to maintain silencing in *h2a.w-2* mutants. To test this hypothesis, we combined *h2a.w-2* with mutants impaired in the deposition of H3K9me (*suvh4/5/6, cmt3*, or *cmt2/3*) or H1.1 and H1.2 (*h1*), which are the predominant linker histone H1 proteins in heterochromatin of vegetative cells (Rutowicz et al., 2019).

Amongst mutants defective in a single chromatin component, the highest number of upregulated TEs was observed in *suvh4/5/6* (**fig 1A**), which comprised most of the TEs upregulated in *cmt3* and *cmt2/3* (**extended fig 1A**). Most of the comparatively lower number of TEs upregulated in *h1* were also upregulated in *suvh4/5/6* (**extended fig 1A**). Additional loss of H2A.W dramatically increased the number of upregulated TEs far above the sum of TEs deregulated in the individual mutant backgrounds, demonstrating a synergistic effect of H2A.W with either H3K9me or H1 on TE silencing (**fig 1A**). A comparable synergy was also observed on the transcript levels of upregulated TEs (**extended fig 1B**).

**Figure 1.**
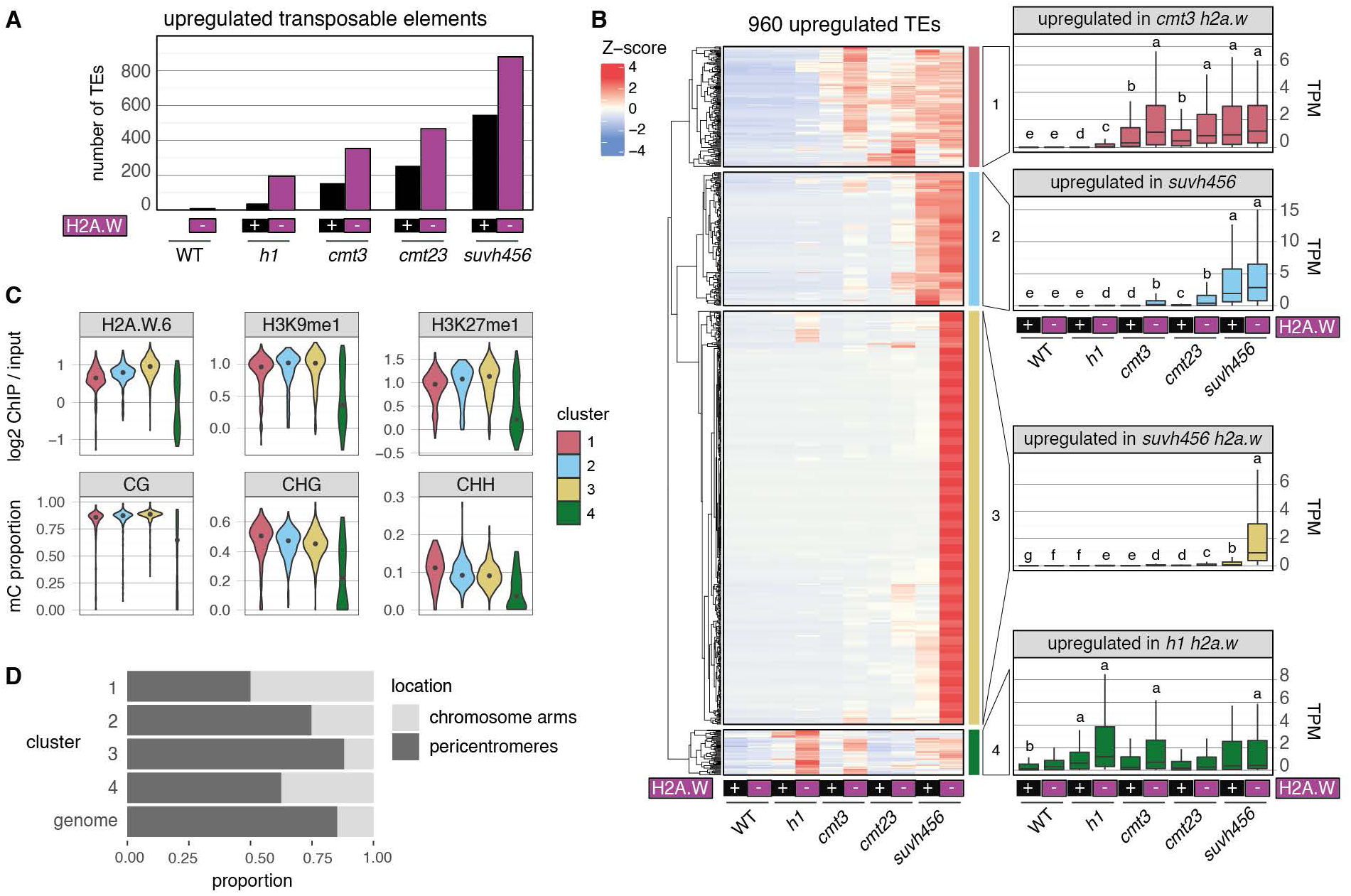
Effects of *h2a.w-2* mutations on release of transposable element silencing in different epigenetic mutant backgrounds. **(A)** Number of upregulated transposable element genes (TEs) in the indicated mutant backgrounds, detected by RNA-seq. **(B)** K-means clustering of all TEs detected as upregulated in the dataset, based on Z-score normalization of transcript per million (TPM) values in the indicated mutant backgrounds. Boxplots show the TPM values for each cluster. Lowercase letters show significant differences between groups evaluated with a Dwass-Steel-Crichtlow-Fligner test (P < 0.05). For clarity, only some groups are displayed for cluster 4. P-values are available in supplementary table 1. (C) Violin plots showing the distribution of wild-type heterochromatin marks in clusters defined in **(B)**, with median represented as black points. **(D)** Compartmentalization of transposable elements in clusters defined in **(B)** between chromosome arms and pericentromeres.

### Highly heterochromatic TEs are kept silent by the combined actions of H3K9 and H2A.W

To characterize the function of H2A.W in silencing TEs in conjunction with heterochromatic features, we selected all TEs upregulated in at least one genetic background and looked at their transcript levels in all genotypes. We used hierarchical clustering to identify dominant trends (**fig 1B**), and determined the enrichment of heterochromatic marks at TEs for each cluster, using previously published wild type datasets (Bourguet et al., 2021; Lorković et al., 2017) (**fig 1C**). We also determined the genomic position of TEs as either pericentromeric or on chromosomes arms, (**fig 1D**) and the distribution of TE superfamilies (**extended fig 1D**).

Cluster 4 comprised 61 TEs upregulated by the joint loss of H2A.W and H1 (**fig 1B**). These TEs showed the lowest levels of heterochromatic marks (DNA methylation, heterochromatin-associated histone modifications and variants) and, surprisingly, the lowest levels of H1 (**fig 1C, extended fig 1C**). One third of the TEs from Cluster 4 localized to chromosomes arms and included a large proportion of LTR Copia elements. H2A.W and H1 synergistically silenced only a subset of the TEs repressed by H2A.W and the other tested pathways, showing that H1 and H2A.W maintain silencing only at a specific set of TEs.

Although transcripts in Cluster 2 (corresponding to 180 TEs) were hardly detected in the absence of CMT2/3, they were upregulated upon additional loss of H2A.W, showing that H2A.W contributes to silencing of these TEs in synergy with CMT2/3. Transcript levels were highest in the absence of SUVH4/5/6, with no significant impact on further loss of H2A.W, indicating that these TEs are primarily silenced by H3K9me (**fig 1B**). This cluster showed a strong enrichment for DNA En-Spm elements and intermediate levels of heterochromatic marks (**fig 1C, extended fig 1C-D**).

Cluster 1 comprised 162 TEs that were synergistically upregulated when *h2a.w-2* mutations were combined with *cmt3* or *cmt2/3* but, in contrast to cluster 2, transcript levels did not increase further in the absence of SUVH4/5/6. Since CMT2/3 deposit CHG and CHH methylation and are directed by SUVH4/5/6 (Stroud et al., 2014), this indicates that TEs in Cluster 1 are controlled by CHG and CHH methylation with a contribution of H2A.W. Accordingly, these TEs showed the highest levels of CHG and CHH methylation (**fig 1C**). TEs in Cluster 1 were mostly present in chromosome arms and predominantly LTR Copia elements (**fig 1D and extended fig 1D**). Such over-representation of LTR Copia elements was reported for regions that lose mCHH in *cmt3* and TEs upregulated in *drm1 drm2 cmt2/3* (Stroud et al., 2013, 2014). This shows that Cluster 1 TEs are silenced by H2A.W and H3K9me-targeted CHG and CHH methylation.

Cluster 3 was the largest and included 557 TEs strongly upregulated only in the absence of both H2A.W and the H3K9 methyltransferases. Interestingly, the inability of *suvh4/5/6* mutations to affect the transcript levels of these TEs correlated with a higher enrichment of H2A.W.6, H3K9me1, H3K27me1, and mCG relative to other clusters (**fig 1C**). This suggests that high levels of other heterochromatic marks could compensate for the loss of H3K9me and CHG and CHH methylation at these elements. They had lower levels of CHG and CHH methylation compared to Clusters 1 and 2 and, accordingly, the loss of CMT2/3 and H2A.W had little impact on their transcript levels. These TEs were present primarily in pericentromeres and were enriched for DNA MuDR and En-Spm elements (**fig 1D and extended fig 1D**).

Overall, our data reveal that TEs are silenced by H2A.W acting together with H1, CH (CHG and CHH) DNA methylation or H3K9me. These pathways affect TEs differently, and these differences are associated with the TE’s superfamily, location, and other chromatin marks. The roles of H3K9me and H2A.W in jointly maintaining transcriptional silencing in Cluster 3 affect 58% of the TEs upregulated in our dataset, most notably pericentromeric TEs with high levels of heterochromatic marks. The much smaller impact of CMT2/3 and H2A.W in the repression of these TEs suggests that the repression is directly mediated by H3K9me and H2A.W, independently of CH methylation.

### H2A.W and the RNA-directed DNA methylation pathway silence different TEs

To further study the role of H2A.W in silencing, we wanted to use mutant combinations generated with the *h2a.w.6-1* mutant allele. However, this allele has undergone genomic rearrangements (Bourguet et al., 2021) that could possibly confound our analysis. For example, we detected 13 upregulated TEs in *h2a.w.6-1/7*, different from the 3 upregulated TEs in *h2a.w-2* (**extended fig 2A**). Therefore, we verified the effect of *h2a.w.6-1/7* in the *h1, cmt2*, and *suvh4* mutant backgrounds, and found that *h2a.w.6-1/7* dramatically increased the number of upregulated TEs in all cases (**extended fig 2B**), similar to the effect observed with the *h2a.w-2* allele (**fig 1A**). This indicates that the loss of H2A.W.6 and H2A.W.7 outweighs any confounding effects of the genomic rearrangements. Therefore, *h2a.w.6-1/7* also are suitable mutations to study the role of H2A.W in silencing. Furthermore, because H2A.W.12 is knocked-out in *h2a.w-2* but not in *h2a.w.6-1/7*, this data indicates that H2A.W.12 doesn’t play a major role in TE silencing, consistent with this gene being lowly transcribed in vegetative tissues (Yelagandula et al., 2014).

**Extended figure 1.**
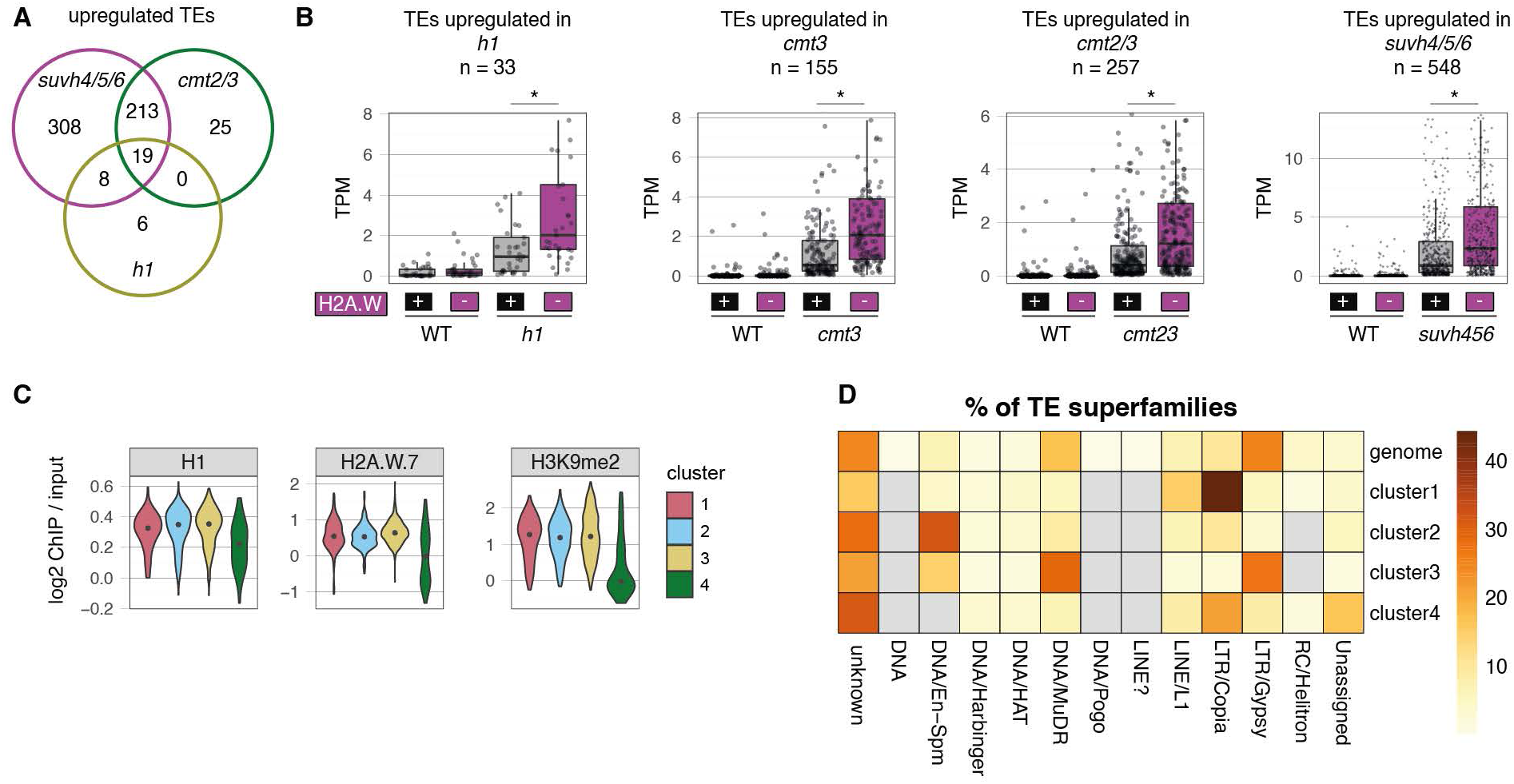
**(A)** Intersect between upregulated transposable element genes (TEs) in the indicated mutant backgrounds, detected by RNA-seq. **(B)** Boxplots show the TPM values of upregulated TEs in the indicated mutant backgrounds. Data points are displayed but outliers exceeding the Y scale upper limit are not displayed. Asterisks indicate statistically significant differences between groups using a two-sided unpaired Wilcoxon rank-sum test (*P* < 0.005). **(C)** Violin plots showing the distribution of heterochromatin marks in wild-type using the clusters defined in **(B)**, with median represented as black points. **(D)** Distribution of TE superfamilies in clusters defined in figure 1B. The percentage of TEs found in a superfamily is shown relative to the total number of TEs in each cluster. Missing data is shown in grey.

H3K9 methylation directs *de novo* methylation mediated by the RdDM pathway (Law et al., 2013; Zhang et al., 2013). RdDM is required to establish *de novo* silencing of certain transgenes (Chan et al., 2004; Kanno et al., 2005; You et al., 2013) and new insertions of the ÉVADÉ transposon (Marí-Ordóñez et al., 2013), but also contributes to silencing maintenance at certain genes and TEs (Chan et al., 2006; Ito et al., 2011; Stroud et al., 2014). RdDM and CMT2/3 are mutually exclusive pathways, showing little overlap (Stroud et al., 2014). For these reasons, we evaluated whether H2A.W cooperates with RdDM to maintain TE silencing. Because H2A.W.12 is hardly expressed in seedlings (Yelagandula et al., 2014), we combined the *h2a.w.6-1/h2a.w.7* double mutant (*h2a.w.6-1/7*) with null mutant alleles of *NRPD2A*, the common subunit of the RNA polymerases IV and V or *DOMAINS REARRANGED METHYLTRANSFERASE 2* (*DRM2*), the final effector of RdDM.

Only a few TEs were upregulated in *drm2* and *nrpd2a* (**fig 2A**), in accordance with previous studies showing a small contribution of RdDM to silencing maintenance at TEs (Stroud et al., 2014). TEs upregulated in *nrpd2a* did not intersect with those upregulated in *h2a.w.6-1/7* (**fig 2B**), suggesting that TEs upregulated in *h2a.w.6-1/7* are not silenced by RdDM. We evaluated the interaction of H2A.W with the RdDM pathway in silencing by looking at synergies between *h2a.w.6-1/7* and either *nrpd2a* or *drm2* mutations. The number of upregulated TEs did not increase in double mutants compared with single mutants (**fig 2A**). Furthermore, expression levels of TEs upregulated in *nrpd2a* and *drm2* did not increase upon further loss of H2A.W (**fig 2C**), in contrast with other mutations (**extended fig 1B**), showing that H2A.W does not contribute to silencing these TEs. The lack of synergy between H2A.W and RdDM mutations in these analyses indicates that these pathways do not cooperate in TE silencing.

**Figure 2.**
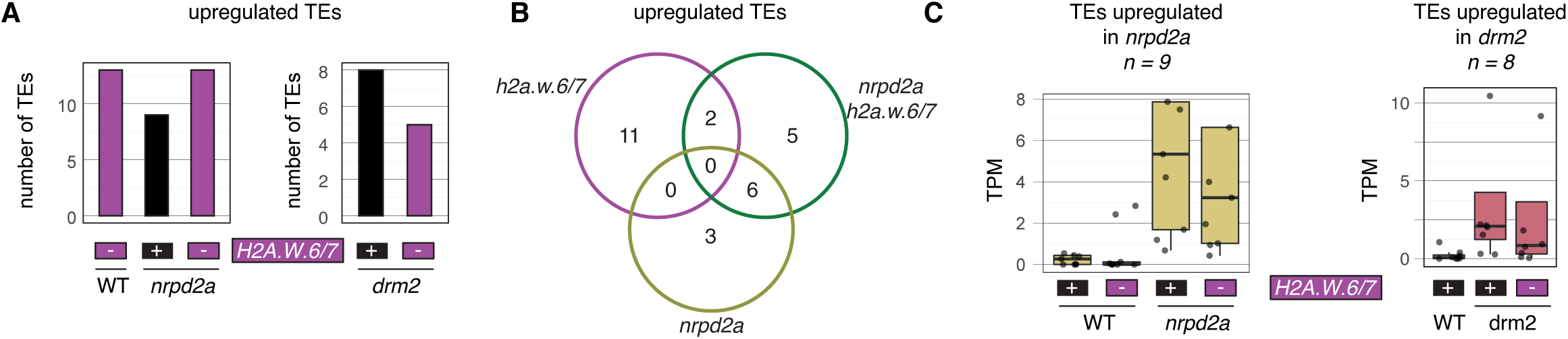
Interaction of the RNA-directed DNA methylation pathway with H2A.W on transposable element transcription. **(A)** Number of upregulated transposable element genes (TEs) in the indicated mutant backgrounds, detected by RNA-seq. **(B)** Overlap between TEs upregulated in the indicated mutants. (**C**)Transcript accumulation at TEs upregulated in *nrpd2a* (left) and *drm2* (right), in WT, *nrpd2a* and *drm2*, with or without *h2a.w.6-1/7* mutations. Transcript per million (TPM) values of RNA-seq read count are shown.

**Extended figure 2.**
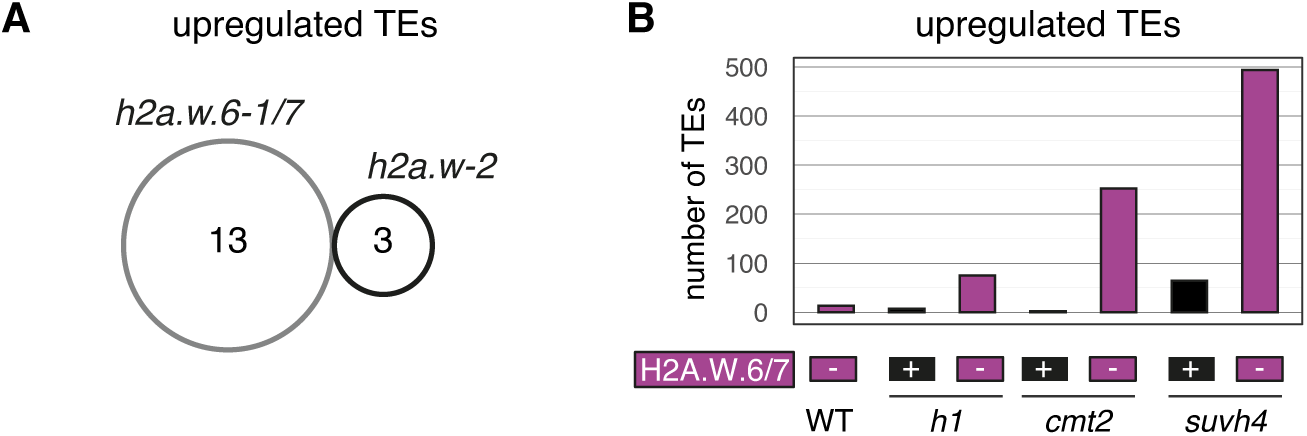
(**A**) Overlap of upregulated transposable element genes (TEs) in the indicated mutant backgrounds, detected by RNA-seq. (**B**) Number of significantly upregulated TEs in the indicated mutant backgrounds.

### Half of DDM1-targeted TEs are silenced by H2A.W and H3K9me

We have previously proposed that the deposition of H2A.W by DDM1 is involved in transcriptional silencing of TEs (Osakabe et al., 2021). To test this model, we evaluated the genetic interaction between *ddm1* and *h2a.w-2*. Relative to the *ddm1* mutant, the number of upregulated TEs or the extent of their upregulation were not increased in *ddm1 h2a.w-2* (**fig 3A, B**). This indicates that DDM1 acts upstream of H2A.W in the maintenance of TE silencing, supporting the model that DDM1 deposits H2A.W.

**Figure 3.**
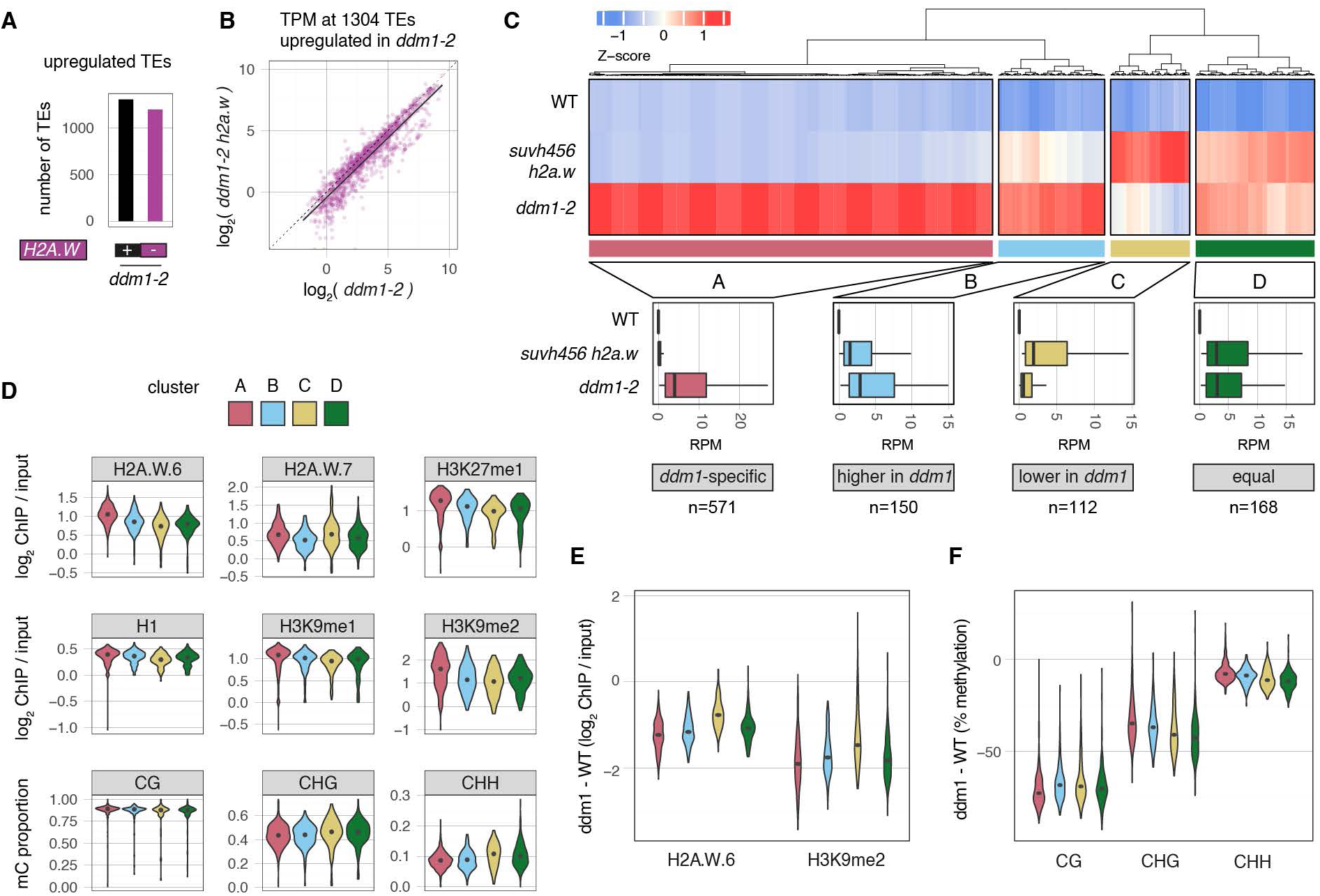
Comparing release of silencing in *suvh456 h2a.w-2* and *ddm1-2* mutants **(A)** Number of upregulated transposable element genes (TEs) in the indicated mutant backgrounds, detected by RNA-seq. **(B)** Transcript accumulation at TEs upregulated in *ddmt-2*, with *ddmt-2* on the x-axis and *ddmt-2 h2a.w-2* on the y-axis. The log_2_ of transcript per million (TPM) values of RNA-seq read count are shown with a regression line and its 95% confidence intervals in black. **(C)** K-means clustering of all TEs detected as upregulated in the indicated mutants, based on Z-score normalization of read per million (RPM) values from QuantSeq libraries of the indicated mutant backgrounds. Boxplots show the RPM values for each cluster. **(D, E, F)** Violin plots showing the distribution of wild-type heterochromatin marks **(D)**, the difference between *ddm1* and WT for the indicated chromatin features **(E)** or methylation context **(F)** at TEs clustered in **(C)**. Dots show the median values.

In *ddm1* mutants, massive TE transcription correlates with a decrease in H2A.W, H3K9me, H3K27me1, and DNA methylation (Ikeda et al., 2017; Osakabe et al., 2021; Vongs et al., 1993). This pervasive and complex alteration of the chromatin landscape makes it difficult to determine which DDM1-dependent marks contribute to silencing and to what extent. To assess the role of H2A.W and H3K9me in DDM1-dependent silencing, we compared the impact of *ddm1* and *suvh4/5/6 h2a.w-2* on TE expression. Nearly all (88%) of the TEs upregulated in *suvh4/5/6 h2a.w-2* were upregulated in *ddm1* (**extended fig 3A**), showing that DDM1 silences TEs by depositing H2A.W and maintaining H3K9me, confirming predictions of DDM1’s role in silencing (Gendrel et al., 2002; Osakabe et al., 2021). Conversely, 55% of *ddm1*-upregulated TEs were not upregulated in *suvh4/5/6 h2a.w-2*, indicating that DDM1 also silences TEs by interacting with other chromatin components.

### Highly heterochromatic TEs silenced by DDM1 are resistant to the concomitant loss of H3K9me and H2A.W

Unsupervised clustering of upregulated TEs revealed four distinct clusters (**fig 3C**). Cluster C comprised 112 TEs with strong upregulation in *suvh4/5/6 h2a.w-2* but no or only low upregulation in *ddm1* mutants, and whose chromatin accessibility was not affected by *ddm1* mutations (**extended fig 3C**). These TEs were characterized by having the lowest enrichment of H2A.W.6, H3K27me1, H1, and H3K9me (**fig 3D**). We speculated that ectopic H3K27me3 deposition in *ddm1* could prevent their upregulation (Rougée et al., 2021), but the increase in H3K27me3 on TEs in cluster C was not higher than other TE clusters (**extended fig 3B**). Changes in DNA methylation did not differ from other clusters (**Fig 3F**) however, TEs in cluster C lost less H2A.W.6 and H3K9me2 (**fig 3E**). Therefore, we conclude that these TEs remain silent in *ddm1* due to retention of H2A.W and H3K9me2, possibly maintained by alternative pathways.

In contrast, chromatin accessibility increased markedly around TEs from Clusters A, B, and D in *ddm1* (**extended fig 3C**). Circa 50% of the TEs upregulated in our dataset were only upregulated in *ddm1* (Cluster A) and were characterized by the strongest enrichment in all heterochromatic histone modifications and H2A.W (**fig 3D**). These TEs are almost exclusively located in pericentromeres and comprise predominantly LTR gypsy elements (**extended Fig 3E**). Cluster B comprised 150 TEs that were more strongly upregulated in *ddm1* than in *suvh4/5/6 h2a.w-2* and cluster D comprised TEs equally upregulated in both mutants (**fig 3C**). Notable differences in the relative enrichment of different TE superfamilies and genomic localization marked each cluster (**extended fig 3E and D**), showing variable degrees of involvement of H3K9me and H2A.W deposition by DDM1 in different regions of the genome. Interestingly, WT levels of H3K9me and H2A.W across clusters followed trends opposite with levels of mCHG and mCHH, suggesting that H3K9me and H2A.W limit chromatin accessibility to the methyltransferases CMT2/3. In contrast, the loss of H2A.W and H3K9me in *ddm1* mutants for all clusters further supports the conclusion that these chromatin components work synergistically to repress TEs. Overall, our data show that H2A.W and H3K9me are responsible for silencing roughly 40% of the TEs upregulated in the *ddm1* mutant. The remaining 60% of the DDM1-controlled TEs are highly heterochromatic Gypsy elements located deep in pericentromeres and remain silent even upon loss of H3K9me and H2A.W in *suvh4/5/6 h2a.w-2*. Chromatin at these TEs is strongly enriched in H3K27me1, H1, and mCG, which could maintain silencing at these TEs in the absence of DDM1.

**Extended figure 3.**
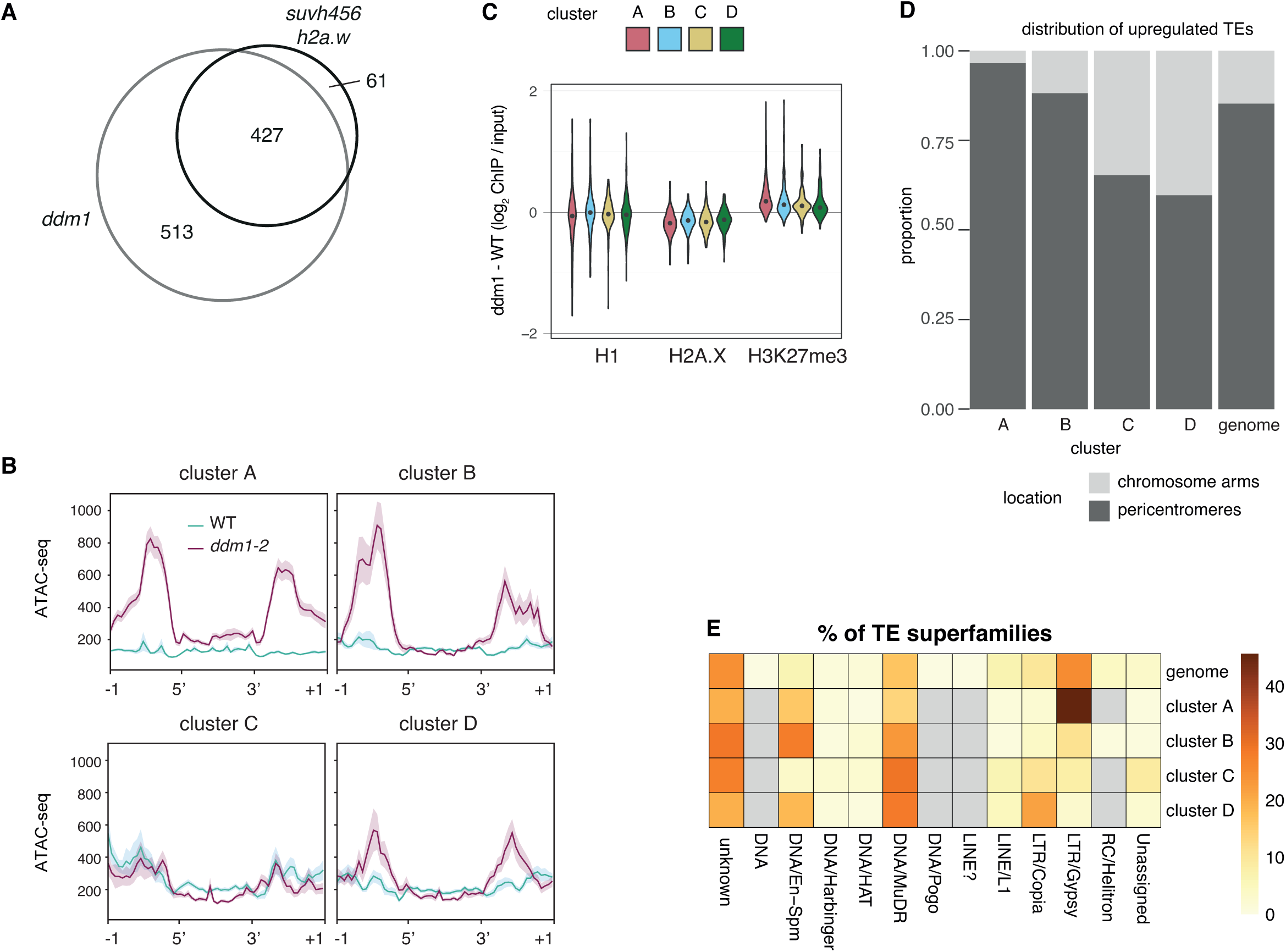
(**A**) Overlap of upregulated transposable element genes (TEs) in the indicated mutant backgrounds, detected by RNA-seq. (**B**) Chromatin accessibility in WT and *ddm1-2* in the clusters defined in figure 3A. Values show the average between replicates of ATAC-seq processed files from GSE155503 (Zhong et al. 2021). (**C**) Violin plots showing the distribution of the difference between ddm1 and WT for the indicated chromatin features in the clusters defined in figure 3A. Dots show the median values. (**D**) Compartmentalization of transposable elements in clusters defined in figure 3A between chromosome arms and pericentromeres. (**E**) Distribution of TE superfamilies in clusters defined in figure 3A. The percentage of TEs found in a superfamily is shown relative to the total number of TEs in each cluster. Missing data is shown in grey.

## Discussion

From these data, we conclude that DDM1 silences TEs by acting upstream of H2A.W in a common genetic pathway. This supports its role in the deposition of H2A.W at TEs in the Arabidopsis genome (Osakabe et al., 2021). The absence of H2A.W alone does not impact TE silencing, in contrast to *ddm1*, because *h2a.w* mutations hardly affect DNA methylation, H3K9me, and H3K27me1 (Bourguet et al., 2021) when compared with the dramatic impact of *ddm1* (Gendrel et al., 2002; Ikeda et al., 2017; Vongs et al., 1993; Zemach et al., 2013). H2A.W’s role in TE silencing is only revealed upon the loss of an additional component of constitutive heterochromatin. Thus, H2A.W acts in synergy with the linker histone H1 and H3K9me to silence TEs that are targeted by DDM1. H2A.W does not cooperate with the RdDM pathway which, remarkably, has been shown to have genomic targets distinct from DDM1-dependent sites (Zemach et al., 2013). Because H2A.W confers specific properties to nucleosome arrays (Osakabe et al., 2018; Yelagandula et al., 2014) and restricts chromatin accessibility (Bourguet et al., 2021), we propose that H2A.W acts synergistically with H3K9me to restrain chromatin accessibility, in a manner comparable to that shown for H2A.W and H1 (Bourguet et al., 2021).

Since many TEs silenced by DDM1 are not upregulated in mutants lacking both H2A.W and H3K9me, other DDM1-dependent heterochromatin components must act to maintain TE silencing, possibly in synergy with H2A.W. Immediate candidates are CG methylation and H3K27me1, since they are lost in *ddm1* mutants (Ikeda et al., 2017; Zemach et al., 2013), but other lesser-known modifications associated with heterochromatin could be involved, such as H4K20me1 (Roudier et al., 2011) and H3K23me1 (Trejo-Arellano et al., 2017). The enzymes responsible for the maintenance of H4K20me1 and H3K23me1 are still unknown and it is thus difficult to assess their impact directly. Here, we partially reproduce TE upregulation in the absence of DDM1 by removing H3K9me and H2A.W, but all heterochromatic marks that co-exist at a single TE locus are likely to contribute to transcriptional silencing (Rigal & Mathieu, 2011). Whether these marks affect the ability of H2A.W nucleosome arrays to prevent RNA Pol II recruitment and transcription remains to be investigated.

In conclusion, we propose that H2A.W interacts with H3K9me to further enhance compaction of nucleosome arrays. The mechanism causing the synergy remains unknown but could rely on phase separation of nucleosomes with H2A.W and H3K9me, since the H3K9me reader protein AGDP1 promotes heterochromatin formation and phase-separation of nucleosome arrays (C. Zhang et al., 2018; Zhao et al., 2018). To maintain DNA methylation at these inaccessible regions, methyltransferases require DDM1 (Lyons & Zilberman, 2017; Zemach et al., 2013). It is possible that the interaction between H3K9me and H2A.W prevents the dynamic exchange of H2A.W and H1, in a manner that would extend previous models proposed to explain the *ddm1* mutant phenotype (Lyons & Zilberman, 2017; Zemach et al., 2013). DDM1 would permit access to DNA methyltransferases but not to the transcriptional machinery, maintaining TEs in a silent state.

The human and murine homologs of DDM1 are Helicase Lymphoid Specific (HELLS) and Lymphoid Specific Helicase (LSH), respectively (Chen et al., 2022). LSH is associated with heterochromatin, and LSH depletion induces DNA hypomethylation, alters chromatin accessibility, and changes histone modifications associated with repressed chromatin (Ren et al., 2015, 2019; Yu et al., 2014). Like DDM1, LSH represses TE transcription (Dunican et al., 2013). LSH binds and deposits the heterodimer macroH2A-H2B in an ATPase dependent manner (Ni et al., 2020; Ni & Muegge, 2021). The variant macroH2A shares some features with H2A.W (Berger et al., 2022); it occupies constitutive heterochromatin (Douet et al., 2017) and is associated with TE silencing (Sun & Bernstein, 2019) but to a much lesser extent than LSH. Therefore, we speculate that joint synergies between macroH2A and heterochromatin modifications play an important role in silencing TE activity in mammals, as shown in Arabidopsis in this study.

Broadly, our study illustrates that histone H2A variants associate with specific epigenetic marks to regulate transcription in a distinct compartment of chromatin. In Arabidopsis, repressive arrays of H2A.Z associate with H3K27me3 and cover repressed genes of facultative heterochromatin (Carter et al., 2018). *In vitro*, arrays of H2A.Z are the preferred substrate of PRC2 (Wang et al., 2018), suggesting a synergistic impact of H2A.Z and H3K27me3 on transcription that remains to be demonstrated *in planta*. H2A.Z and PRC2 are deeply conserved features of heterochromatin in Eukaryotes (Grau-Bové et al., 2022), while H2A.W evolved at a later stage (Lei et al., 2021). Thus, the impact of H2A.W in cooperation with heterochromatic marks demonstrated here likely illustrates a general and ancient phenomenon in eukaryotic chromatin, that associates H2A variants with chromatin marks to control transcription.

## Methods

### Plant material and growth conditions

All mutants are in the Columbia-0 (Col) ecotype. h2a.w-2, h2a.w.6-1 (SALK_024544.32) and h2a.w.7 (GK_149G05) mutant lines were described previously (Bourguet et al., 2021; Yelagandula et al., 2014). h1.1 (SALK_128430C), h1.2 (GABI_406H11), cmt2-3 (SALK_012874), cmt3-11 (SALK_148381) and suvh4/5/6 triple mutant (suvh4: SALK_044606, suvh5: SALK_207725, suvh6: SAIL_1244_F04) were described previously (Ebbs & Bender, 2006; To et al., 2020; Zemach et al., 2013). The EMS-induced SNP in ddm1-2 has been described (Vongs et al., 1993). drm2-2 (SALK_150863), nrpd2a-1 (SALK_095689), cmt3-11 (SALK_148381) and kyp-6 (SALK_041474) mutants were used in combination with h2a.w.6-1/7. Arabidopsis plants were grown at 21°C under long day conditions (16 hours light and 8 hours dark).

### Library preparation for RNA-sequencing

Seedlings were grown vertically in half Murashige and Skoog medium in MES buffer with 1% Plant Agar. Total RNA was extracted from ten-day-old whole seedlings (roots and aerial parts) with RNeasy Mini kit (Qiagen, Chatsworth, USA). RNA was subjected to a DNase treatment using DNase Free Kit (Ambion, Austin, USA).

Poly-A libraries (figure 1 and related data) were constructed with the NEBNext Ultra RNA library prep kit for Illumina (New England Biolabs, Ipswich, USA), following the manufacturer’s instructions. 75 bp paired-end reads were generated on an Illumina NextSeq 550 system. Quantseq libraries (figure 3C) were prepared using QuantSeq 3’ mRNA-Seq Library Prep Kit (FWD) for Illumina (Lexogen, Vienna, Austria), following the manufacturer’s instructions. 75 bp single-end reads were generated on an Illumina NextSeq 550 system. Libraries depleted of ribosomal RNA (figure 2 and 3A-B) were constructed with the TruSeq Total RNA with Ribo Zero plant Kit (Illumina, San Diego, USA), according to the manufacturer’s instructions. We sequenced 100 bp single-end reads on an Illumina Hiseq 2500 system (figure 2) or 50 bp single-end reads on Hiseq v4 system (figure 3A-B).

### RNA-sequencing analysis

Reads from poly-A and Ribo-Zero libraries were trimmed using Trim Galore (version 0.6.2) and default parameters. QuantSeq reads were trimmed to remove adapter and polyA sequences using bbduk from bbmap (version 38.18) following Lexogen’s recommended parameters (k=13 ktrim=r useshortkmers=t mink=5 qtrim=t trimq=10 minlength=20). From there, reads from all kind of library were processed identically as indicated below.

Reads were aligned using STAR (version 2.7.1a) to the TAIR10 genome allowing 4% mismatches, using annotations based on the Araport11 assembly. Reads mapping at multiple locations were kept, considering all alignments with the best mapping score as primary alignments if they map to 20 or less locations (--bamRemoveDuplicatesType UniqueIdenticalNotMulti --outSAMprimaryFlag AllBestScore --outFilterMultimapNmax 20). Reads were counted using featureCounts from the Subread package (version 2.0.1). Multimapping reads were given a value of 1 / *n*, where *n* is the number of possible mapping loci (-s 1 -O -M --fraction --primary - Q 0).

After rounding up gene counts, significantly differentially expressed genes were selected using DESeq2 (Love et al., 2014) with a minimum absolute log2 fold-change of 1 and an adjusted *p-value* equal or below 0.05. Genes covered with less than 10 reads in the considered dataset were discarded beforehand.

Read counts were normalized by transcript per million (TPM) for ribo-depleted and Poly-A libraries. For Quantseq libraries, read counts should not be normalized by gene length since only the 3’ end of transcripts is sequenced, hence we used read per million normalization (RPM). The average between biological replicates was subsequently used to represent transcript levels. Heatmaps were plotted with the R package ComplexHeatmap (version 2.6.2), using both k-mean clustering (consensus of 1000 runs) and unsupervised row clustering with the “euclidean” mode.

### Annotations and pericentromeric coordinates

We restricted the analysis to the 3901 transposable elements annotated as *transposable_element_gene* in the Araport11 GFF annotation reference (https://www.arabidopsis.org/download_files/Genes/Araport11_genome_release/Araport11_GFF3_genes_transposons.May2022.gff.gz). Pericentromeres and chromosome arms were distinguished based on H3K9me2 profiles, as defined previously (Bernatavichute et al., 2008).

### Quantifying DNA methylation, histone marks, histone variants and accessibility using previously published datasets

We used deeptools (version 3.3.1) *multiBigwigSummary* to compute the enrichment of BS-seq, ChIP-seq or ATAC-seq reads over a given set of TEs, starting from processed data (bigwig files) available from previously published studies. The average between replicates was computed with deeptools’ *bigwigCompare* tool. For WT levels of DNA methylation, H3K9me1, H3K9me2, H3K27me1, H1, the data originated from GSE146948 (Bourguet et al., 2021). For WT levels of H2A.W.6 and H2A.W.7, we used GSE95557 (Lorković et al., 2017). ATAC-seq data for WT and *ddm1* mutants came from GSE155503 (Zhong et al., 2021). The effect of *ddm1* mutations on DNA methylation, histone marks and histone variants was calculated using data from GSE150436 (Osakabe et al., 2021). For metaplots, we used *deeptools computeMatrix scale-regions*, fitting regions of interest to 1000-bp and plotting 1000-bp upstream and downstream using a window size of 50-bp.

### Statistical analysis

Analyses were conducted in the R environment (version 4.0.3) (R Core Team, 2019). Boxplots followed Tukey’s definition, with whiskers extending to the furthest point that is less than 1.5-fold the interquartile range from the box. We used the native *krukal.test* R function to perform Kruskal-Wallis rank-sum test, and Dwass-Steel-Critchlow-Fligner post hoc tests were made using the pSDCFlig function with the asymptotic method from the NSM3 package (Schneider et al., 2018). Other statistical tests were performed with the corresponding native R functions.

## Acknowledgements

We thank James Watson for suggestions and critical reading of the manuscript. We would like to thank the Plant Sciences and Next Generation Sequencing at the Vienna BioCenter Core Facilities (VBCF), and the Molecular Biology Service core facilities. This work was funded by FWF grants I2203, P26887, P32054, and P33380 to F.B., M2539-B21 to A.O., the Japan Society for the Promotion of Science (JSPS) Overseas Research Fellowships (to A.O.), JST, PREST (JPMJPR20K3 to A.O.) and the Human Frontier Science Program RGP0025/2021 to T.K.. This project has received funding from the European Union’s Framework Programme for Research and Innovation Horizon 2020 (2014-2020) under the Marie Curie Skłodowska Grant Agreement Nr. 847548 to P.B., from the Japanese Ministry of Education, Culture, Sports, Science and Technology 22K06180, 19H05740 and 17K15059 to T.K.T., 21K20628 to A.O., 19H00995 and 21H04977 to T.K., and from the Ministry of Science and Technology of Taiwan 104-2923-B-001-003-MY2 and 109-2313-B-001-009-MY3 to P-Y.C.

## Author contributions

P.B., R.Y., P-Y. C., and F.B. conceived and designed the experiments. R.Y., A.O. and T.K.T. generated the genetic material used in this study. P.B., R.Y. and A.O. performed the RNA-seq measurements. A.A. and R.J.L. performed the first round of bioinformatic analyses. P.B. performed the bioinformatic and statistical analyses and curated data presented in the manuscript. F.B., P-Y. C., and T.K. supervised the study. P.B, F.B. and R.Y. wrote the manuscript with input from all other authors.

## Competing interest declaration

The authors declare no competing interests.

**Correspondence and requests for materials** should be addressed to Frédéric Berger.

